# A Dual-Pathway Wnt-IL-13 Fusion Protein Enhances Human Intestinal Regeneration Through Tuft Cell Activation

**DOI:** 10.64898/2026.01.31.702996

**Authors:** Li Yang, Lulu Huang, Fenna L.H. van Rijt, Baojie Zhang, Amir Giladi, Jochem H. Bernink, Hans Clevers, Claudia Y. Janda

## Abstract

Compromised intestinal regenerative responses drive severe inflammatory conditions, such as inflammatory bowel disease and graft-versus-host disease, affecting millions of people worldwide each year. Despite extensive research, effective therapies remain limited, and no curative treatments are currently available. We recently discovered that human intestinal tuft cells promote tissue repair following injury through Wnt and IL-4/IL-13 signaling pathways. Building on this discovery, here we report the engineering and functional validation of a synthetic Wnt-IL-13 fusion protein that simultaneously activates both the Wnt and IL-4/IL-13 signaling pathways to enhance human intestinal tuft cell activity. Employing human organoids technology, we demonstrate that this therapeutic approach promotes mucosal healing.

## Introduction

Continuous intestinal epithelial renewal is essential to maintain or restore epithelial integrity in case of injury, preventing bacterial dissemination and uncontrolled inflammation. This process is orchestrated primarily by intestinal stem cells (ISCs) and transient amplifying cells, which continuously replenish the epithelium^1,2^. A coordinated network of growth factors and morphogens acting at the bottom of intestinal crypts on ISCs that preserve stemness and drive continuous epithelial proliferation^3-5^. Disruption or loss of ISCs delays epithelial renewal and compromises barrier integrity. As a consequence, bacterial translocation into otherwise sterile mucosal tissues can occur, triggering severe inflammatory responses. Such compromised mucosal integrity drives multiple clinical conditions, such as intestinal mucositis in leukemic patients that undergo preconditioning regimen prior receiving a hematopoietic stem cell graft^6^, or severe intestinal ulcers in patients that suffer from inflammatory bowel disease (IBD)^7^. While disease severity in such patients is assessed by various endoscope-based scoring systems that center on mucosal healing, current treatment options for intestinal inflammation are primarily aimed at suppressing inflammation rather than boosting the epithelial restorative response itself.

Recent insights into human mucosal regenerative biology have identified tuft cells as a rare epithelial cell type with epithelial reparative properties^8^. Mechanistically, tuft cell development and proliferation critically depend on the Wnt and interleukin-4 (IL-4) / interleukin-13 (IL-13) signaling pathways. Upon activation, tuft cells can act as a local cellular source of growth factors such as Epiregulin (Ereg) and heparin-binding epithelial growth factor (HB-EGF), and can transdifferentiate into ISCs, thereby contributing to tissue regeneration. Given the roles of Wnt and IL-4/IL-13 signaling in promoting intestinal regeneration, simultaneous therapeutic activation represents a promising approach, though it faces several obstacles.

Wnt ligands activate signaling by binding to Frizzled (FZD) and LRP5/6 receptors. The 10 FZDs (FZD1-10) exhibit distinct yet overlapping expression patterns across tissue cell types, though their individual contributions to regenerative responses remain incompletely understood. A major limitation for therapeutic development is that Wnt proteins require lipid modification for receptor binding, rendering them highly hydrophobic and unsuitable for drug development^9,10^. However, the discovery that forced proximity of FZD and LRP5/6 receptors is sufficient to induce Wnt/β-catening signal has enabled the development of antibody-based Wnt surrogates^11-13^. These engineered bispecific ligands promote FZD-LRP5/6 heterodimerization, effectively mimicking natural Wnt ligands while circumventing the challenges of working with native Wnt proteins. As a result, various Wnt surrogates are being developed as potential regenerative therapeutics^14-18^.

Similarly, therapeutic activation of the IL-4/IL-13 axis presents both opportunities and challenges. These cytokines, which are crucial for regulating immune responses and tissue repair, signal through type I and type II receptor complexes involving IL-4 receptor alpha (IL-4Rα) paired with either the common gamma chain (γc), or IL-13 receptor alpha 1 (IL-13Rα1), respectively^19^. While IL-4 can engage both type I and type II receptor complexes, IL-13 is restricted to signaling through the type II receptor complex. The widespread expression of IL-4/IL-13 receptors across diverse cell types underlies their central role in type 2 immune responses, including Th2 cell differentiation, M2 macrophage polarization, and eosinophil recruitment^20^. However, this broad receptor distribution complicates therapeutic applications, as systemic administration risks off-target effects including uncontrolled inflammation, excessive mucus production, and tissue fibrosis. Consequently, strategies that spatially restrict IL-4/IL-13 signaling to the epithelium while minimizing systemic exposure could enhance safety profiles for regenerative applications.

Here, we leverage human intestinal organoid models to explore a novel class of therapeutic strategies aimed at promoting mucosal healing by targeting tuft cells. Specifically, we engineered a fusion protein that simultaneously activates Wnt and IL-13 signaling to boost the regenerative properties of human intestinal tuft cells. We demonstrate a proof-of-concept approach for the development of alternative therapeutics that primarily acts on enhancing epithelial restoration rather than solely suppressing inflammation.

## Results

### Wnt and IL-4/IL-13 receptor expression profiles in tuft cells inform the engineering strategy of Wnt-IL-13 fusion proteins

In a previous study^8^, tuft cells were shown to express high levels of IL-13Rα1 and, to a lesser extent, IL-4Rα in human primary intestinal tissues, making IL-13 the cytokine of choice for our study. Furthermore, its more restricted receptor specificity and reduced pleiotropy confer a narrower and more defined target cell range than IL-4. To design an appropriate engineering strategy for a Wnt–IL-13 fusion protein, we first characterized the expression profiles of Wnt receptors across human intestinal epithelial lineages, with particular focus on tuft cells. Specifically, we profiled the expression of the 10 FZDs and LRP5/LRP6 co-receptors using a previously generated single-cell RNA sequencing (scRNA-seq) dataset derived from ileum organoid-derived epithelial cells cultured in tuft cell differentiation medium, with and without exposure to IL-4/IL-13^8^. This dataset contained four distinct tuft cell subpopulations, in addition to stem cells and two goblet cell types.

Across the four tuft cell subsets, FZD5 and FZD3 were strongly expressed, with moderate expression of FZD6 (Figure 1A–B, Suppl. Figure 1). FZD9 expression was more restricted, predominating in the tuft-1 and tuft-2 subsets. Notably, FZD5, FZD3, and FZD6 were also detected in stem and goblet cells, suggesting broader functions for these receptors beyond the tuft cell lineage. Both LRP5 and LRP6 were consistently expressed across all tuft subsets. To validate these observations, we analyzed an independent scRNA-seq dataset of primary human adult small intestine^21^ (Figure 1C–D, Suppl. Figure 2). The expression pattern in primary tissue was more heterogeneous; however, LRP5 and LRP6 remained strongly expressed in tuft cells. FZD5 again emerged as the predominant FZD receptor, albeit at lower levels compared to the tuft cells in organoids as well as all other cells within the intestinal epithelial lineage, spanning stem cells, transit amplifying cells, enterocytes, goblet cells, and enteroendocrine cells. Taken together, these data thus provide a rationale for developing Wnt-IL-13 fusion proteins that engage FZD5 to drive robust activity or FZD9 to achieve greater selectivity for tuft cells.

**Figure 1.**
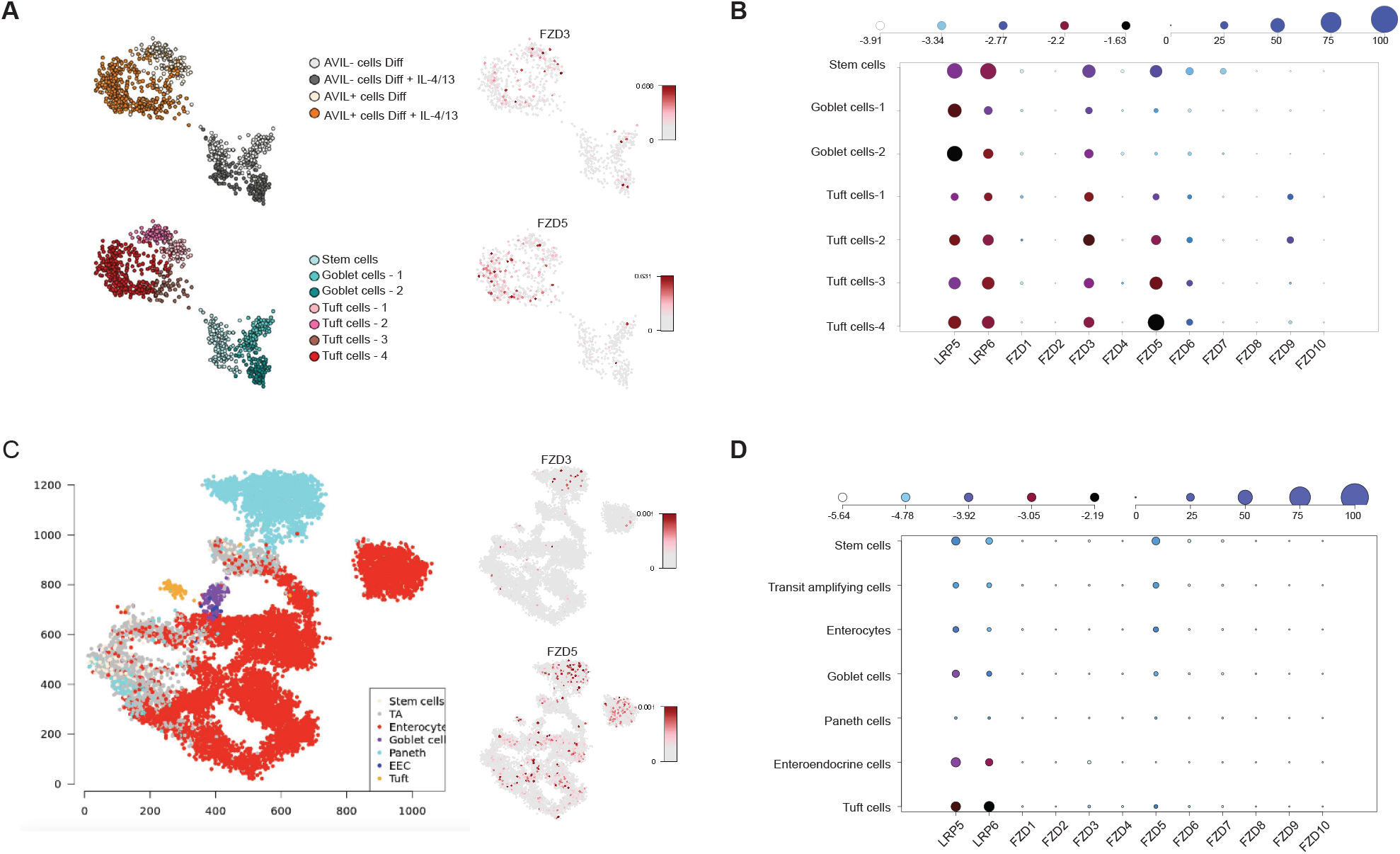
Wnt receptor expression profiles in human intestinal epithelial cells. **(A)** Metacell 2D representation of scRNA-seq data of ileum-derived organoids cultured in differentiation media with and without IL-4/IL-13 activation^8^, n=953 cells colored by their medium conditions, cell subtypes, as well as log-normalized expression of FZD5 and FZD3. **(B)** Expression of the Wnt receptors FZD and LRP5/6 across the scRNA-seq dataset in (A). Dot color relates to size-normalized mean expression values and dot size to fraction of expressing cells. **(C)** Metacell 2D representation of scRNA-seq data of primary human adult small intestine^21^, n=15,184 single epithelial cells colored by their cell subtypes, as well as log-normalized expression of FZD5 and FZD3. **(D)** Expression of the Wnt receptors FZD and LRP5/6 across the scRNA-seq dataset in (C). Dot color relates to size-normalized mean expression values and dot size to fraction of expressing cells.

### Engineering of tri-specific Wnt-IL-13 fusion proteins for the simultaneous activation of Wnt and IL-13 signaling

To test whether tuft cells can be expanded through simultaneous activation of Wnt and IL-13 signaling, we generated a panel of Wnt–IL-13 fusion proteins aimed at robustly and simultaneously activating Wnt and IL-13 signaling within the same cell. These constructs combined previously validated Wnt surrogate formats, comprising anti-FZD antibodies and a single-domain variable domain (VHHs) that binds LRP5/6 fused to the N-terminus of the light chains, with wild-type human IL-13 connected via a flexible peptide linkers (Figure 2A)^13^. We took advantage of publicly available antibodies and antibody fragments. Two anti-FZD antibodies were selected based on their binding to FZD5: mAb1, which binds FZD1/2/5/7/8^22^, and mAb2, which binds FZD1/2/4/5/7/8^23^; and a LRP6-binding VHH^24^.

**Figure 2.**
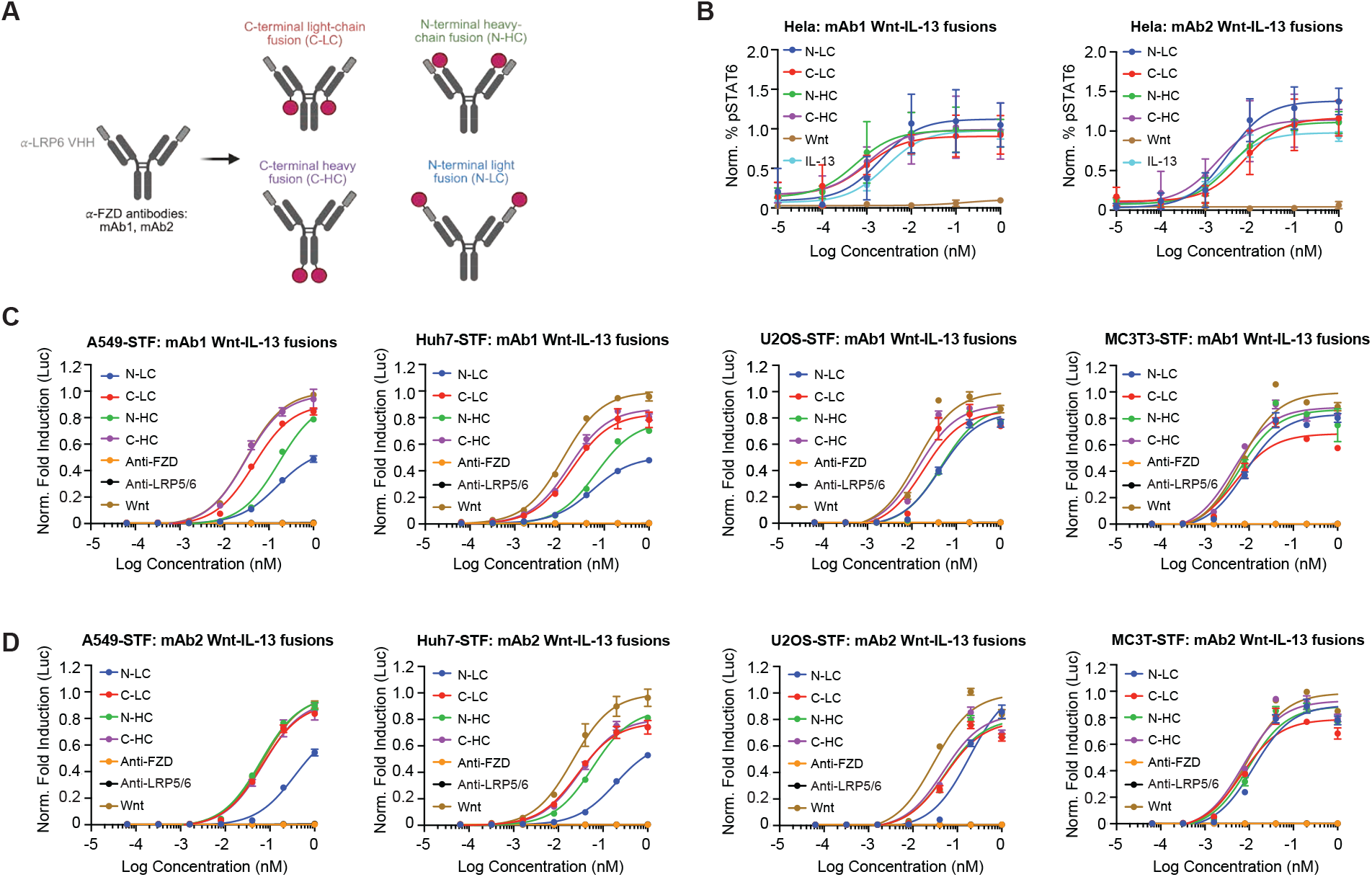
Functional characterization of tri-specific Wnt–IL-13 fusion proteins. **(A)** Schematic representation of the tri-specific Wnt–IL-13 fusion protein formats. The anti-FZD IgG antibodies are shown in black, the anti-LRP5/6 VHH domain in gray, and IL-13 as a red circle. Created with BioRender.com **(B)** STAT6 phosphorylation induced by IL-13 and the Wnt–IL-13 fusion proteins in HeLa cells, measured by FACS. The percentage of pSTAT6-positive cells in stimulated samples minus unstimulated samples was normalized relative to the pSTAT6 level observed with recombinant IL-13. Data represent mean ± SD of three biological replicates, each containing the mean of at least two technical replicates. **(C–D)** Activation of the β-catenin–dependent STF reporter by Wnt surrogates (mAb1 and mAb2) and Wnt–IL-13 fusion proteins in A549 (expresses predominantly FZD2 > FZD6 > FZD7 > FZD3), Huh7 (FZD5 > FZD4 > FZD7 > FZD6 > FZD1), U2OS (FZD6 > FZD3 > FZD1 > FZD7), and MC3T3 cells (FZD1>FZD7), and shown as fold induction of luciferase activity in stimulated relative to unstimulated samples, and normalized to the signal of the Wnt surrogate. Data represent mean ± SD of three biological replicates, each containing the mean of three technical replicates.

Because the addition of IL-13 could sterically interfere with receptor binding by either the Wnt surrogate (to FZD and LRP5/6) or IL-13 (to IL-13Rα1 and IL-4Rα), IL-13 was fused to four different positions, either the N- or C-terminus of the heavy or light chain of the anti-FZD antibody, to identify a geometry that optimizes dual signaling. All constructs, including control Wnt surrogates and antibodies, were transiently expressed in Expi293 cells and purified by Protein A affinity chromatography followed by size-exclusion chromatography.

We first examined whether the fusion proteins retained IL-13 signaling activity by measuring STAT6 phosphorylation in HeLa cells, which express IL-13 receptors but display low basal STAT6 activation. All Wnt–IL-13 formats induced STAT6 phosphorylation with similar potency to recombinant IL-13, independent of fusion orientation, indicating that IL-13 activity was preserved (Figure 2B).

Next, we assessed Wnt signaling activity using the SuperTOPFlash reporter assay, a standard readout of Wnt/β-catenin signaling, in several cell lines (A549, Huh7, U2OS, MC3T3) with distinct FZD expression patterns (Figure 2C-D). In the mAb1 context, C-terminal IL-13 fusions to either the light (C-LC) or heavy chain (C-HC) showed comparable activity to the parental Wnt surrogate, whereas N-terminal fusions (N-LC, N-HC) displayed reduced signaling in A549 and Huh7 cells but comparable activities to the positive control in U2OS or MC3T3 cells. In the mAb2 context, only the Wnt-IL-13 fusion in which IL-13 linked to the N-terminus of the light-chain consistently showed lower activity compared to the Wnt surrogate itself, across cell types. All others (C-LC, C-HC, N-HC) showed comparable activity to the Wnt surrogate.

Together, these data indicate that C-terminal fusions of IL-13 generally preserve Wnt and IL-13 signaling activity more effectively than N-terminal fusions, likely by minimizing steric hindrance between the IL-13 ligand and the Wnt surrogate domains. This panel of engineered Wnt–IL-13 fusion proteins provide a versatile platform to study the coordinated activation of Wnt and IL-13 signaling pathways.

### Wnt-IL-13 fusion protein enhances human intestinal tuft cell regeneration

We previously took advantage of human organoid technology to systematically dissect niche factors and requirements for tuft cell development and proliferation. Building on these insights, we employed our engineered human intestinal tuft cell reporter organoids^8^ to assess the activity of two Wnt-IL-13 fusion proteins, with IL-13 fused to the C-terminus of the light chain of mAb1 and mAb2 (Suppl. Figure 3A). To identify the most promising Wnt-IL-13 fusion protein for activating intestinal tuft cells, we took a stepwise approach. We first compared the activities of the Wnt surrogates based on mAb1 or mAb2 with the commercial Wnt Surrogate-Fc, referred to as NGS Wnt. Relative to base medium lacking Wnt, tuft cell frequency increased approximately 7-fold in the presence of NGS Wnt. The mAb2 and mAb1-based Wnt surrogates showed a 5 and 8-fold increase, respectively, over baseline (Figure 3A). The difference in activity may be related to intrinsic differences of mAb1 versus mAb2, such as affinities, rather than related to the FZD binding specificities of the mAb1/2 themselves. Based on its superior activity, subsequent experiments focused on the mAb1-based Wnt-IL-13 fusion protein with IL-13 fused to the C-terminus of the light chain (C-LC).

**Figure 3:**
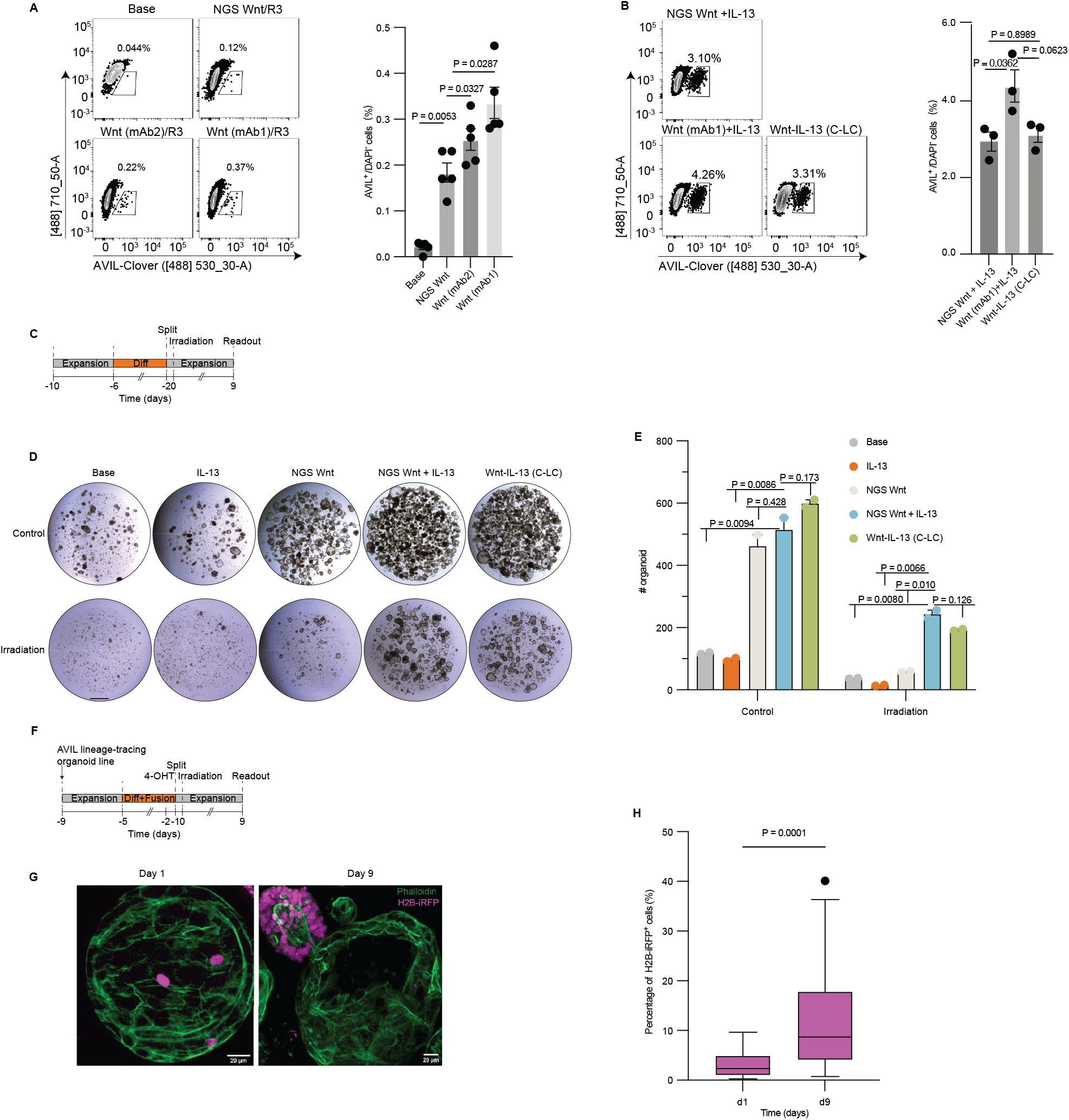
Wnt-IL13 fusion protein in human intestine organoids. **(A-B)** Representative flow cytometry (left) and quantification (right) of the AVIL ^+^ cell percentage of DAPI^−^ cells in human ileum AVIL-Clover reporter organoids differentiated in differentiation regimens. Each dot is a well. **A**) *n* = 5 wells per condition (pooled from three independent experiments). **B**) *n* = 3 wells per condition, results are representative of three independent experiments. **(C–E)** Schematic (**C**), representative images (**D**), quantification of organoid numbers (**E**) from organoids following irradiation. Experiments were performed on two clones (Suppl. Figure 3B). **E**), Each dot is one BME drop. *n* = 2 drops per condition. **(F–H)** Schematic (**F**), representative images (**G**) and quantification (**H**) of AVIL lineage-tracing organoids after irradiation. **H**), Results are pooled from three independent experiments, *n* = 26 (day1) or 32 (day9) organoids. **A**,**B**,**E**, Data are presented as mean ± SEM. **H**, Boxplots show data from the 25th to 75th percentiles and whiskers extending to the minimum and maximum within 1.5× inter-quartile range, with dots marking outliers. **A**,**B**,**E**, P values are derived from *One-way ANOVA with Dunnett’s test is used for multiple comparisons* against the Wnt(UPE)/R3 group (**A**) or *One-way ANOVA with* Tukey’s multiple comparisons test (**B**). **E**, *P* values are derived from false discovery rate (FDR)-adjusted unpaired two-tailed Student’s *t*-test against the fusion protein group. Scale bars, 1 mm (**D**) and 20 µm (**G**). R3, R-spondin3.

Consistent with prior findings that the addition of recombinant IL-13 promotes tuft cell proliferation^8^, reporter organoids cultured with NGS Wnt plus IL-13 triggered a 20-fold increase in tuft cell frequency as compared to NGS Wnt alone. This effect was even more pronounced by the mAb1-based Wnt surrogate plus IL-13 (Figure 3B). Notably, tuft cell reporter organoids cultured with the Wnt-IL-13 (C-LC) expanded comparably to those treated with IL-13 plus mAb1-based Wnt surrogate, demonstrating the potency of a single synthetic protein to induce tuft cell expansion in human intestinal organoids (Figure 3B).

To test whether Wnt-IL-13 (C-LC) promotes tuft cell-mediated regeneration in a setting of epithelial injury, we irradiated differentiated human intestinal tuft cell reporter organoids, and monitored recovery over a 9-day period (Figure 3C). Organoids treated with NGS Wnt or IL-13 alone failed to recover, whereas organoids exposed to either combined compounds or Wnt-IL-13 (C-LC) successfully regained growth capacity (Figure 3D-E, Suppl. Figure 3B). To corroborate these findings, we performed experiments with tuft cell lineage-tracing organoids, subjected to irradiation and recovery (Figure 3F). Wnt-IL13 (C-LC) treatment restored organoid growth and resulted in tuft cell-derived organoid segments (iRFP^+^ cells), along with an overall increase in iRFP-labelled cell frequency (Figure 3G-H).

Together, these findings demonstrate that the engineered Wnt-IL-13 fusion protein potently activates tuft cells and induces a regenerative response in compromised human intestinal epithelium *in vitro*.

## Discussion

In this study, we leveraged human intestinal organoid technology to demonstrate that human intestinal tuft cells are therapeutically targetable epithelial cells with potent regenerative capacity. We show that a synthetic fusion protein with drug-like properties that co-activates Wnt and IL-13 signaling enhances the regenerative response of the human intestine after radiation injury by engaging tuft cell–mediated repair. These findings build on our previous work demonstrating that loss of canonical ISCs can be compensated by activation of tuft cells, a rare epithelial population positioned at the crypt–villus junction, and that their reparative function can be amplified through concurrent activation of Wnt and IL-13 signaling. Together, these results provide the first translational demonstration that targeted activation of tuft cell–driven repair can be harnessed to restore mucosal integrity, for instance in inflammatory and cytotoxic injury settings.

Current therapies for the management of intestinal inflammatory conditions that emerge as a consequence of severe mucosal barrier disruption primary aim on suppressing the inflammatory cascade, while a focus on adequately promoting mucosal healing is lacking. Our work demonstrates that selective boosting tuft cell activity may provide a complementary and potentially synergistic strategy that directly enhances epithelial repair. The Wnt-IL-13 fusion protein described here may therefore serve as a prototype for therapies designed to restore, rather than merely preserve, intestinal function, by simultaneously activating the Wnt and IL-13 signaling pathways, without the need for providing the individual agonists exogenously. Nevertheless, our approach has several limitations and will benefit from further refinement. These considerations align with broader gaps in the translation pipeline for IL-13 and Wnt-based therapies and are mostly related to their inherent biochemical properties and pleiotropic activities.

The clinical translation of IL-13 and IL-4 agonists as therapeutic agents remains largely unexplored. Most insights stem from preclinical models investigating signaling through the type II receptor complex (IL-4Rα/IL-13Rα1), which is predominantly expressed on non-hematopoietic cells such as epithelial, stromal, and tumor cells^20^. In these models, IL-4 and IL-13 have been shown to promote wound healing and tissue repair, highlighting opportunities in regenerative medicine. However, their dual nature poses major clinical challenges, as the same pathways that mediate tissue regeneration also drive pathological fibrosis and allergic inflammation. Engineered cytokine variants, including IL-4 superkines and IL-13 analogs with altered receptor-binding specificity, offer a way to harness their beneficial effects while limiting adverse outcomes^25,26^. In particular, IL-13 variants with increased affinity for the IL-13Rα1 receptor chain present a promising avenue for further exploration in the context of the Wnt-IL-13 fusions. Moreover, the clinical development of cytokines therapeutics faces additional hurdles, most notably suboptimal pharmacokinetic and stability profiles, challenges that are actively being addressed through the design of the Wnt-IL-13 fusion proteins.

Furthermore, the natural Wnt ligands represent powerful modulators of tissue repair and regeneration and the development of Wnt surrogate technology has addressed the limitation of the unfavorable developability of Wnt ligands as therapeutic agents by demonstrating that forced heterodimerization of the Wnt receptors FZDs and LRP5/6 is sufficient to activate β-catenin– dependent Wnt signaling, thereby bypassing the need for the natural lipidated ligand^11,12^. Several Wnt surrogates have since been developed for systemic delivery, showing efficacy in promoting tissue regeneration and repair in preclinical models of bone (osteoporosis, non-union fracture), intestine, and liver injury^13^. Despite this progress, clinical translation remains challenging. The only reported clinical trial involving systemic administration of an antibody-based Wnt surrogate for inflammatory bowel disease was discontinued after phase I due to elevated liver enzyme levels, suggestive of potential hepatotoxicity. Hence, future work on optimizing the activity of the Wnt-IL-13 fusion protein could explore strategies to confine its activity specifically to the intestine. One avenue could involve engineering tetra-specific variants that additionally target tuft cells by incorporating a binding domain for a tuft cell–specific receptor. Alternatively, scRNA-seq data from human intestinal tissue indicate that FZD9 is most differentially expressed in tuft cells. FZD9 displays a more restricted expression pattern compared to the broadly expressed receptors FZD1/2/5/7/8. Notably, while most current anti-FZD antibodies target FZD1/2/5/7/8, redirecting specificity toward FZD9 could further enhance selectivity for tuft cells and minimize off-target effects.

## Material and Methods

### Expression and purification of Wnt-IL13 variants and control antibodies

The coding sequences of the Wnt–IL-13 variants, anti-FZD antibodies, and anti-VHH–Fc variants (for sequences, see Appendix A) were cloned into the pcDNA3.1 vector containing an N-terminal signal peptide. Proteins were expressed by transient transfection of Expi293F cells (Gibco™; Thermo Fisher Scientific) using the ExpiFectamine™ 293 Transfection Kit (Gibco™; Thermo Fisher Scientific), following the manufacturer’s instructions. After four days of expression, the conditioned media were clarified by centrifugation, and the secreted proteins were captured using CaptivA™ Protein A affinity resin (Repligen). Bound proteins were eluted with IgG elution buffer (Thermo Fisher Scientific) containing 300 mM NaCl and immediately neutralized with 100 mM Tris (pH 8.0) to reach a final pH of 7.0. Subsequently, proteins were further purified by size-exclusion chromatography (SEC) on a Superdex 200 Increase 10/300 GL column (Cytiva) using an ÄKTA Pure chromatography system (Cytiva). The running buffer consisted of 1× HBS (20 mM HEPES, pH 7.3; 300 mM NaCl). Purified proteins were concentrated using Amicon® Ultra centrifugal filters (Millipore; MWCO 30 kDa) and sterilized by filtration through 0.22 µm centrifugal filters (Merck). Protein concentrations were determined by NanoDrop A280 measurements using the respective molecular weights and extinction coefficients.

### Cell lines for measuring signaling activities

A549, Huh7, MC3T3, and U2OS cell lines, stably transfected with the Wnt-responsive STF reporter plasmid, were a kind gift from the laboratory of Prof. Dr. K. Christopher Garcia (Stanford University). HeLa cells were purchased from ATCC. A549 cells were cultured in RPMI-1640 medium (Gibco); Huh7 and HeLa cells in high-glucose DMEM (Gibco); MC3T3 cells in α-MEM (Gibco); and U2OS cells in McCoy’s 5A medium (Gibco), each supplemented with 10% fetal bovine serum (Sigma), 1% penicillin– streptomycin (Gibco), and 1% GlutaMAX (Gibco). All cells were maintained in a humidified incubator at 37 °C with 5% CO_2_ and passaged using 0.25% trypsin–EDTA (Gibco) at a split ratio of 1:4 to 1:16. All cell lines were routinely tested for mycoplasma contamination by the mycoplasma testing team at the Princess Máxima Center.

### SuperTop Flash (STF) assay to measure Wnt activity

WNT/β-catenin signaling activity of the Wnt-IL-13 variants was measured in cell lines (A549, Huh7, MC3T3, and U2OS) containing a stably transfected luciferase reporter gene under the control of a WNT-responsive promoter (SuperTop Flash reporter - STF). Cells were seeded in 100 µL of culture medium at a density of 1 × 10^5^ cells per well in flat-bottom 96-well plates and incubated overnight at 37 °C with 5% CO_2_. The following day, Wnt-IL-13 variants and the Wnt surrogate without IL-13 fusion (positive control) were added at concentrations ranging from 1.0 nM to 0.164 pM (5-fold serial dilutions) to stimulate WNT signaling overnight at 37 °C with 5% CO_2_. Anti-FZD antibodies and an anti-LRP5/6 VHH-Fc were included as negative controls. After 20 hours of stimulation, the medium was removed, and cells were lysed in 40 µL of 1× lysis buffer. From each well, 20 µL of lysate was transferred to a white flat-bottom 96-well plate, followed by the addition of 20 µL of luciferase substrate (Promega). Luminescence was measured using a SpectraMax iD3 plate reader (Molecular Devices). The fold induction was calculated according to formula: *Fold Induction* = *Luminescence*_*treated*_/ *Luminescence*_*untreated*_.). Assays were performed in technical and biological triplicates, and results are presented as the mean ± standard deviation of the normalized values of the biological triplicates in relation to the activity of the Wnt surrogate without IL-13. Dose–response curves were generated in GraphPad Prism 10 and nonlinear regression analysis was applied for curve fitting.

### STAT6 phosphorylation assay to measure IL-13 activity

IL-13 signaling activity of the Wnt-IL-13 fusion proteins was assessed in a dose-response assay on HeLa cell lines by measuring STAT6 phosphorylation (pSTAT6). HeLa cells (1.25 × 10^5^) were seeded in 500 µL high-glucose complete DMEM medium (Gibco) in flat-bottom 24-well plates and incubated overnight at 37 °C in 5% CO_2_. The next day, Wnt-IL-13 fusion proteins at different concentrations (0.001 nM to 100 nM, 5-fold serial dilutions) were added to the cells to stimulate IL-13 signaling for 20 minutes. Subsequently, the cells were washed with PBS, dissociated, collected, and permeabilized with ice-cold methanol (Boom, 100%, v/v) at 4 °C for 15 minutes. After removing the methanol, the cells were washed, resuspended in FACS buffer, and transferred to V-bottom 96-well plates for staining. Anti-human pSTAT6-PE (CST) and anti-human STAT6-Alexa Fluor 488 (R&D Systems) antibodies were used at a 1:50 dilution and cells were stained for 45 minutes at 4 °C. Cells were analyzed by FACs using a Beckman Coulter CytoFLEX S (Beckman Coulter, Brea, CA, USA). The extent of STAT6 phosphorylation was determined by subtracting the % of pSTAT6 cells in the unstimulated samples from that of the stimulated samples. Assays were performed in technical and biological triplicates, and the results are presented as the mean ± standard deviation of the normalized values of the biological triplicates in relation to the activity of recombinant IL-13. Dose–response curves were generated in GraphPad Prism 10 and nonlinear regression analysis was applied for curve fitting.

### Tuft cell differentiation in human ileum organoid

Human ileum AVIL-Clover tuft cell reporter and AVIL-lineage tracing organoids were established and cultured as described in^27^. Briefly, organoids were split once a week by mechanical dissociation and cultured in expansion medium as previously described^27^. To differentiate tuft cells, human ileum organoids were expanded for 4 days in expansion medium, then organoids were washed for 30 minutes in DMEM+++: advanced Dulbecco’s modified Eagle’s medium/F12 (Gibco) supplemented with 100 U ml^−1^ penicillin–streptomycin (Gibco), 10 mM HEPES (Gibco), 1× Glutamax (Gibco), and the medium was replaced for tuft cell differentiation medium: 0.5 nM NGS Wnt surrogate (IPA) or 0.5 nM mAb1/mAb2-based Wnt surrogates, 20% (v/v) R-spondin1 (conditioned medium), 10 μM Notch inhibitor DAPT (Sigma-Aldrich), 1× B-27 Supplement (Life Technologies), 1.25 mM *N*-acetylcysteine (Sigma-Aldrich) and 1% (v/v) recombinant Noggin (IPA). In some experiments, 0.5nM human IL-13 (Promega), or 0.5nM Wnt-IL-13 fusion protein were used in this study.

For irradiation of AVIL lineage-tracing organoids (Figure 3F-I), organoids were first differentiated for 4 days in tuft cell differentiation medium containing Wnt-IL-13 fusion protein. They were then treated with 1 μM tamoxifen for 20 hours, followed by splitting and irradiation (9 Gy) one day post-splitting.

### Irradiation in organoids

Culture plates were sealed airtight and irradiated with a single dose of 6 Gy (Figure 3C-E) or 9 Gy (Figure 3F-I) using a linear accelerator (Elekta Precise Linear Accelerator 11F49, Elekta). Plates were placed on a 2 cm-thick polystyrene base and submerged in a 37 °C water bath. Irradiation was delivered from below, with the plates positioned exactly 100 cm from the radiation source. Medium was refreshed after irradiation.

### Flow cytometry

Organoids were dissociated into single cells using TrypLE (TrypLE Express, Life Technologies) with 10 μM Rho kinase inhibitor (Abmole) in 37 °C and mechanical disruption by pipetting every 5 minutes. Then cells were stained with DAPI (4′,6-diamidino-2-phenylindole) and visualized using a BD LSR Fortessa X20 4 laser (BD Biosciences, FACSDiva v.9.0) based on fluorescence levels. FlowJo software (v.10.6.2) was used to analyze the flow cytometry data.

### Whole-mount staining of organoids

Organoids were removed from the BME, then were fixed for 30 minutes at room temperature in formalin. Next, the organoids were permeabilized using 0.1% Tween 20 (Sigma-Aldrich) in PBS for at least 15 minutes and blocked for at least 1 hour in 0.1% Triton X-100 (Sigma-Aldrich), 1 g l^−1^ bovine serum albumin (Sigma-Aldrich) in PBS. Organoids were incubated with Alexa Fluor 488 phalloidin (1:1,000, Thermo Fisher Scientific, A12379) in blocking buffer containing DAPI (1:1,000, Invitrogen, D1306). Sections were embedded in fructose–glycerol clearing solution, then visualized on Leica SP8 confocal microscope (LAS X v.1.1). Image analysis was performed using ImageJ (Fiji, v.1.51n) software.

## Acknowledgements

L.Y. and L.H. acknowledge financial support from the China Scholarship Council program (grant no. 202308440170) and (grant no. 201906210081), respectively. J.H.B. acknowledges financial support from ZonMw Veni fellowship (grant no. 09.150.161.810.107) and Dutch Lung Fund grant (no. 4.2.18.237). The work was further supported by KiKa core funding (C.Y.J.), and Oncode Accelerator, a Dutch National Growth Fund project under grant number NGFOP2201 (C.Y.J). We thank Fleur van der Sterren for technical assistance, the Flow Cytometry Facility of the Princess Máxima Center for their assistance with FACS experiments and the Faculty of Veterinary Medicine of the Utrecht University for supporting the irradiation experiments.

## Authors contributions

L.Y. designed and performed experiments, interpreted results. L.H. designed and performed experiments, interpreted results, and wrote the paper. F.L.H.v.R. designed and performed experiments. B.Z. design and performed experiments. A.G. interpreted results and analyzed data. J.H.B. conceptualized and supervised the project and wrote the paper. H.C. conceptualized and supervised the project and wrote the paper. C.Y.J. conceptualized and supervised the project and wrote the paper.

## Data availability

To support the main finding of this paper, we re-analyzed single-cell RNA-seq data from the following sources: Ileum-derived human organoids under Wnt and IL-13 signaling (<u>GSE233451</u>, Fig. 1A-B), and the Gut Cell Atlas (https://www.gutcellatlas.org/), 10.5061/dryad.8pk0p2ns8, and used for Fig. 1C-D. Source data are provided with this paper.

## Declaration of generative AI usage

In the final stages of preparing this manuscript the authors used Grammarly to improve language and grammar. The authors reviewed and edited the content as needed and take full responsibility for the final manuscript.

---

**Supplementary Figure 1:**
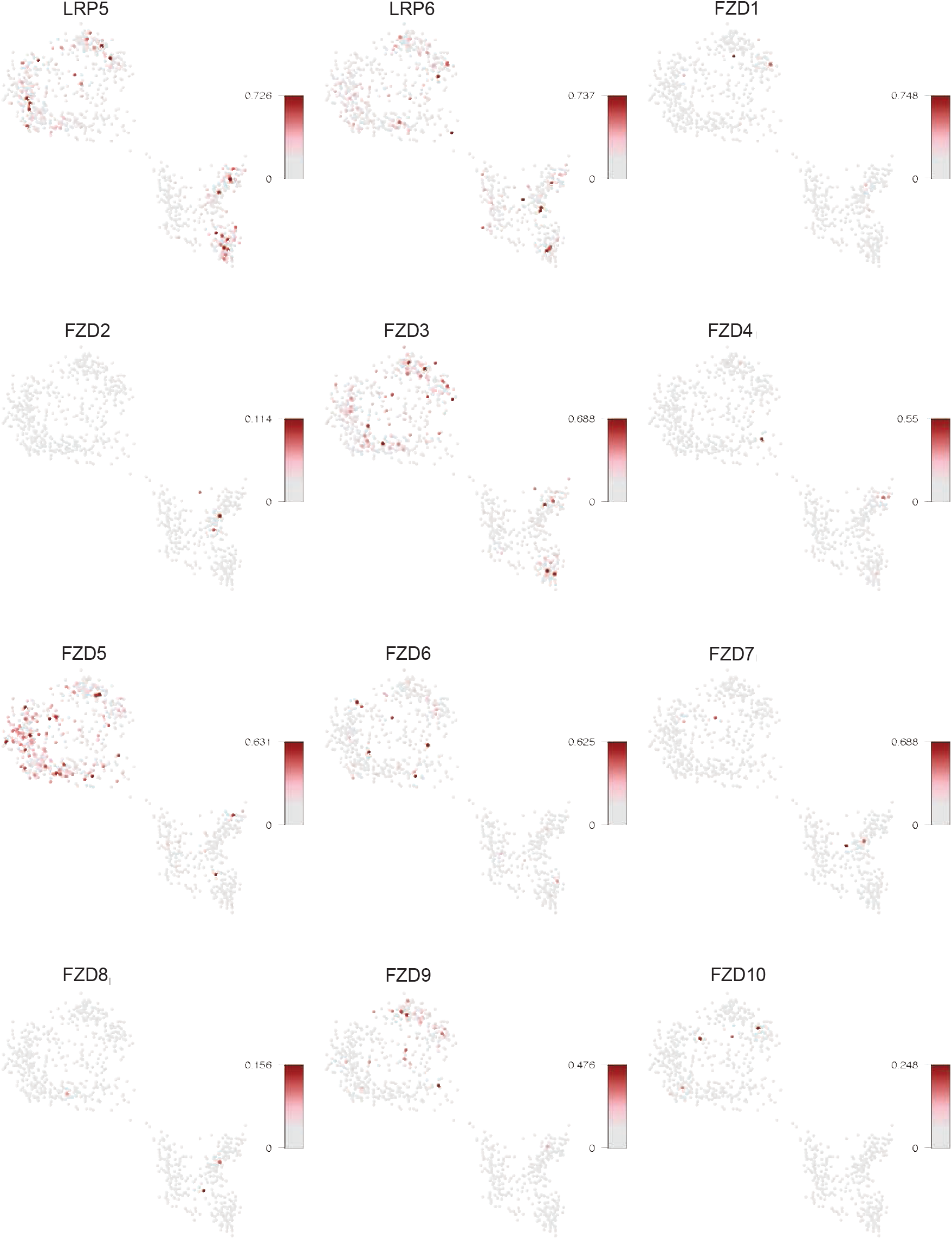
FZD and LRP5/6 expression in ileum-derived organoids. Log-normalized expression of the ten FZDs and LRP5/6, projected on a Metacell 2D representation of scRNA-seq data of ileum-derived organoids cultured in differentiation media with and without IL-4/IL-13 activation^8^.

**Supplementary Figure 2:**
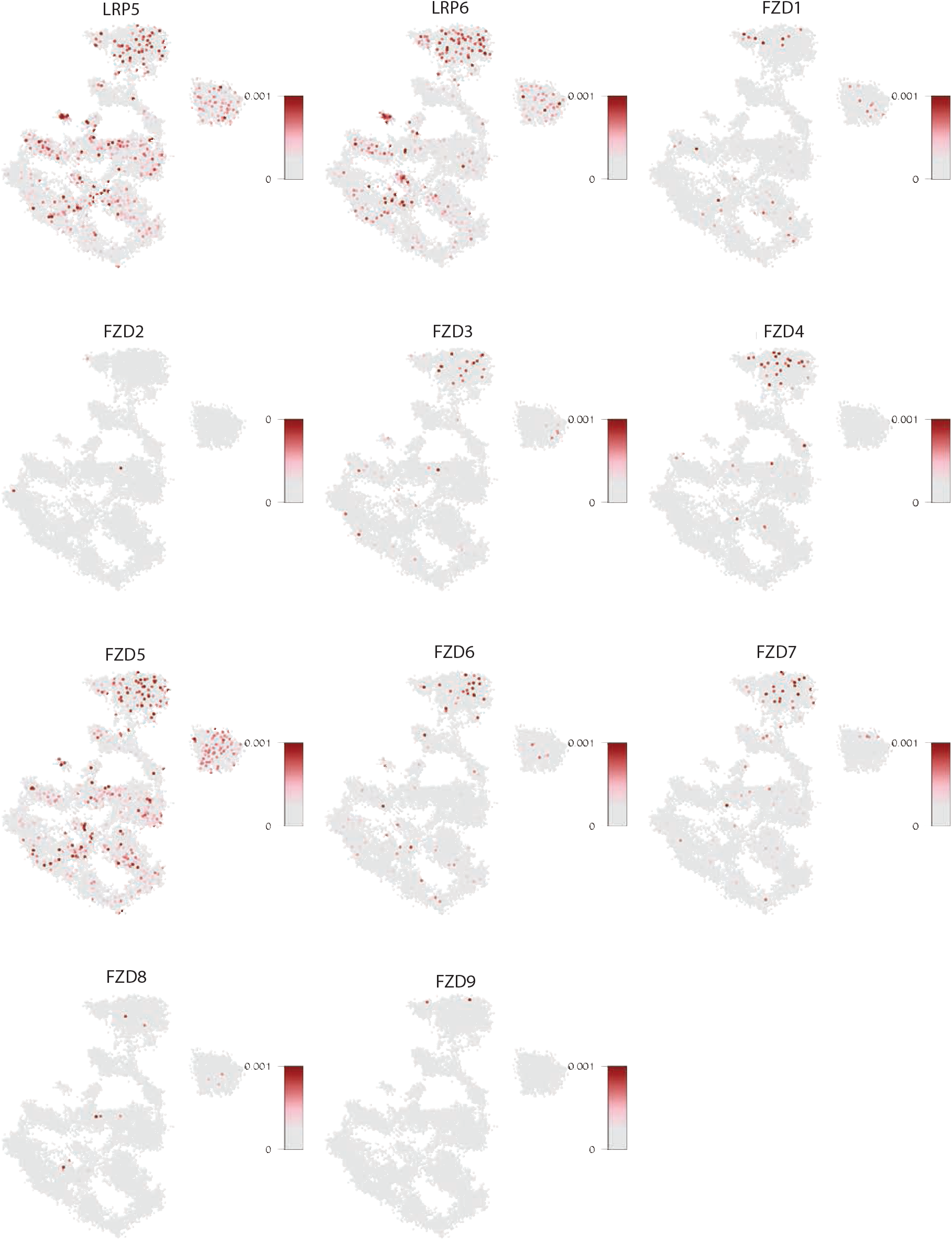
FZD and LRP5/6 expression in human adult small intestine. Log-normalized expression of the ten FZDs and LRP5/6, projected on a Metacell 2D representation of scRNA-seq data of primary human adult small intestine^21^.

**Supplementary Figure 3:**
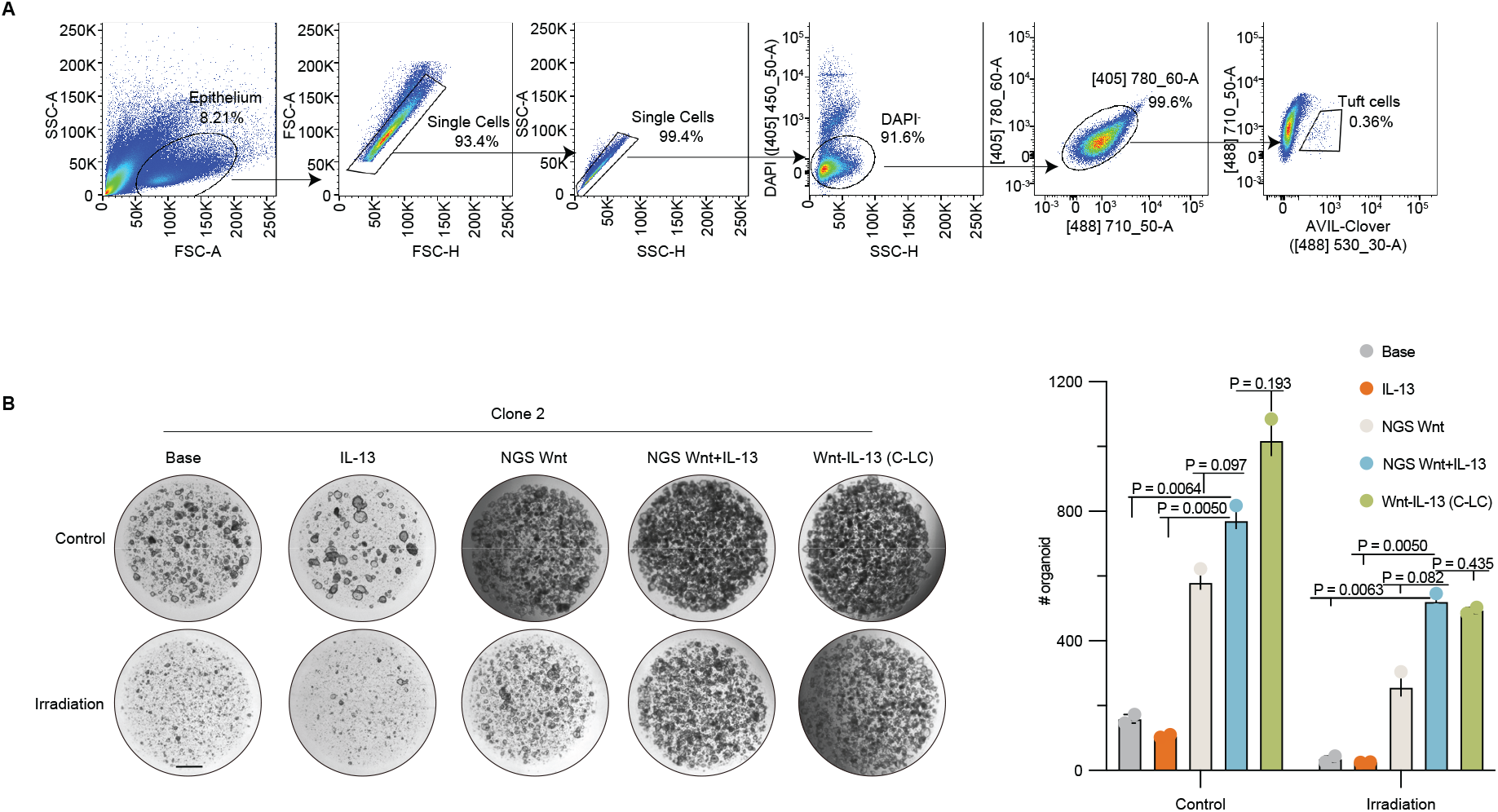
Wnt-IL-13 fusion proteins in human intestine organoids. **(A)** Gating strategy for flow cytometric analysis of AVIL-Clover reporter organoids. **(B)** Representative images of the second organoids cultured in different regimes following irradiation (related to Figure 3C). Scale bar, 1 mm.

